# cadmus: a robust pipeline for scalable retrieval of full-text biomedical literature

**DOI:** 10.64898/2026.05.16.725623

**Authors:** Jamie Campbell, Antoine D. Lain, T. Ian Simpson

**Author notes:** These authors contributed equally (Jamie Campbell, Antoine D. Lain).

## Abstract

`cadmus` is an open-source Python toolkit for automated retrieval and processing of full-text biomedical literature. It utilises programmatic access to PubMed, Crossref, Europe PMC, PMC, and publisher APIs, allowing users to construct large, domain-specific corpora with minimal manual intervention. `cadmus` parses PDF, HTML, XML, and plain text files, standardising them for downstream biomedical text mining. During the retrieval of a Developmental Disorders Corpus (204,043 publications), it achieved an 85.2% full-text retrieval rate with institutional subscriptions and 54.4% without. To test the fidelity of retrieved full-texts, we used ScispaCy to infer the similarity of paired documents from 44,264 open-access PubMed Central files and the files retrieved from `cadmus`, resulting in an average cosine similarity score of 0.98. Rarefaction analyses demonstrated that full-text corpora double the coverage of unique biomedical concepts over abstracts, resulting in better access to the depth of the biomedical information available.

**Availability and implementation:** `cadmus` is a freely available package for non-commercial research at https://github.com/biomedicalinformaticsgroup/cadmus and released under the MIT License.

## Introduction

The biomedical research literature is a vast and rapidly growing scientific resource, yet much of its content remains underutilised for text mining analysis (1, 2). While PubMed provides a comprehensive index of over 39 million bibliographic records, it primarily offers abstracts, which often compress key details about experimental design, methodology, and findings that are only available in the full text. The PubMed Central Open Access (PMC-OA) set currently contains approximately 7 million full-text articles, providing partial access to large-scale text mining. However, it represents only a fraction of the literature indexed in PubMed, leaving many publications inaccessible for biomedical text mining use. Creating large, domain-specific full-text corpora, therefore remains a challenge for Biomedical Natural Language (BioNLP) applications, including knowledge graph construction (3, 4), and reliable biomedical question answering systems (5–7). The difficulties of building a full-text corpus result from a combination of legal, technical, and practical barriers. Publisher websites use heterogeneous formats, ac-cess controls, licensing conditions, and technical inconsistencies in document layouts make parsing error-prone. Existing approaches to biomedical information extraction (8–10) use PubMed abstracts and/or full-text documents from the PMC-Open Access (PMC-OA) collection restricting coverage to only a small section of the published literature. `cadmus` was developed to address these issues by offering an automated, open-source pipeline for programmatic retrieval, parsing, and structuring of biomedical full texts. It leverages existing Application Programming Interfaces (APIs) from data hubs like NCBI PubMed, Crossref, and from individual publishers, but simplifies the requirements for utilising each individual component (11–13), to deliver corpora in a consistent format that is ready for downstream text mining analysis. The system prioritises scalability, fault tolerance, and requires minimal human intervention, making it practical for small targeted queries and the construction of large, domain-specific corpora.

### Software description

The `cadmus` pipeline consists of three sequential stages: ‘query and metadata retrieval’, ‘document acquisition’, and ‘parsing and storage’. The process begins with either a PubMed query string or a list of PMIDs as input, using the Entrez Direct E-utilities service provided by the NCBI to obtain publication metadata (11). Optional API keys for Elsevier and Wiley can be supplied to extend access to full-text retrieval at the second stage. When using Entrez Direct, metadata including titles, abstracts, MeSH terms, author keywords, DOIs, and PMCIDs are collected and stored in structured ‘.zip’ JSON files.

Multiple retrieval strategies are used to fetch articles. `cadmus` first queries the Crossref API (13) to find URLs linked to the article and will follow URL redirection links to find final targets for article source files. These requests can lead to the retrieval of files in XML, HTML, PDF, or plaintext format from several sources per article. Article PMCIDs are also used to query Europe PMC and PMC APIs to fetch open access XML or PDF versions of articles. Where closed source article links are found the ability to retrieve files is dependent on use of the appropriate API key and/or any institutional IP-bounded licencing restrictions. Files are only retrieved if the user has the necessary permission to do so as determined by the relevant licensing conditions for each publisher. `cadmus` uses the word count, file size, and text similarity to the corresponding Entrez Direct abstract to assess whether the correct files have been retrieved. If so, the relevant meta-data is updated in the retrieval catalogue and the process moves on to the next article. This approach minimises the number of API calls and downloads needed per article, identifies and removes erroneous files, and improves the fidelity and efficiency of retrieval. The pipeline includes robust error handling, allowing interrupted processes to be resumed from a saved index as well as allowing for incremental updates by only retrieving publications not already in the corpus.

In the final stage, the retrieved files are parsed using custom scripts that extract and standardise the article text producing clean plain-text files for downstream text-mining applications. For XML and HTML files, `cadmus` uses Beautiful Soup (15), PDFs are processed with Tika (16), and plain-text files are read directly. All files are compressed into per article ‘.zip’ archives, reducing the storage space required and preserving data for future re-use. To simplify data handling, `cadmus`includes a utility function, `parsed_to_df`, that enables users to load article metadata and full-text content directly from these archives without de-compression, supporting storage efficient large-scale corpus exploration.

## Results

`cadmus` was evaluated on a corpus of 204,043 publications generated by combining the PubMed search results for 120 gene names and symbols taken from the Developmental Disorders Genotype-2-Phenotype (DDG2P) dataset (17, 18). With institutional subscriptions, the pipeline successfully retrieved 173,786 full-text documents, achieving an 85.2% retrieval rate. To evaluate retrieval rates without subscription, the same process was run on a Google Colab notebook for 20,000 randomly selected articles from the same corpus. This process retrieved 10,873 articles (54.4%) when running on a private wireless network and 17,040 articles (85.2%) when running on the university network. Processing throughput averaged approximately 250 documents per hour on a single CPU, with memory usage remaining below 4 GB throughout the pipeline. For this specific corpus, PubMed provides a title and abstract text for 179,389 (87.9%) articles, 2.7% more than our full-text corpus. When evaluating the 30,257 articles that were not retrieved as full-text, 16,149 (53.4%) were published by journals that are not covered by the University subscription portfolio, nor did they have a permissive Creative Commons license. To evaluate the fidelity of content retrieved by `cadmus`, we compared 44,264 documents obtained through the pipeline (in all possible formats) with their corresponding versions in the PMC-OA collection. For each document pair, we converted the text into numerical vectors, and we used cosine similarity to measure how closely the two versions matched. Cosine similarity gives a score based on how many words and phrases the documents have in common and how frequently they appear. A score of 1 indicates that the two documents are essentially identical, while a score near 0 shows they are very different. The average score of 0.98 confirms that `cadmus` retrieves accurate copies of full-text files. To confirm our evaluation, we ran the same calculation on random, unrelated document pairs from the corpus, which produced an average similarity of 0.23. The high scores obtained in the matched pairs are due to accurate retrieval rather than chance overlap. Parsing quality was validated by measuring the proportion of out-of-vocabulary (OOV) tokens using ScispaCy (19) found after we parsed the full-text from the original file. These tokens are strings, characters, words or symbols that do not appear in the model vocabulary, often reflecting errors, unusual formatting, or other noise. Text retrieved with `cadmus` had half the OOV rate of the PMC-OA text (0.026 vs 0.049 per token), suggesting that `cadmus` output is cleaner and contains fewer parsing artefacts, making it easier to process for downstream tasks. We evaluated the benefit for information extraction of using article full-text content over abstract text alone by comparing their rarefaction curves for unique Unified Medical Language System (UMLS) Metathesaurus concepts identified using ScispaCy’s entity linker pipeline. Normlalised entity density in full-text documents is comparable to that of abstracts, even though full texts are over 20 times longer indicating, as expected, that concepts are distributed throughout articles (Figure 1A) Unique entity coverage from full-text articles more than doubles the number of unique UMLS concepts compared to abstracts, reflecting substantial information loss when relying only on summaries (Figure 1B). As our test corpus is generated by a query for articles associated with developmental disorders we next evaluated the rarefaction of phenotype concepts by mapping UMLS concepts to the Human Phenotype Ontology (HPO).

**Fig. 1.**
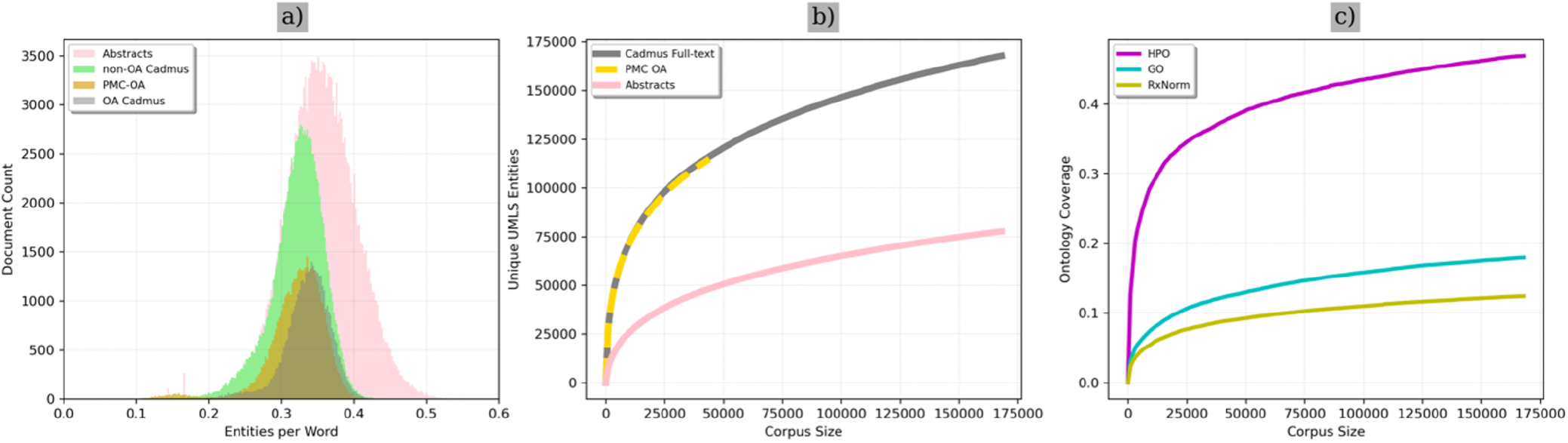
a): Overlapping histograms showing the number of biomedical entities per word for all the documents in each corpus. b): Rarefaction curves showing the total number of unique UMLS concepts (14) found in each corpus, measured at 1000 random document increments. c): Rarefaction curves showing the proportion of each ontology covered by 1000 random document increments for the complete `cadmus` retrieved corpus only. (OA: open access; PMC: PubMed Central; UMLS: Unified Medical Language System; HPO: Human Phenotype Ontology; GO: Gene Ontology; RXNorm: Normalised Naming system for generic and branded drug

### Limitations and Considerations

`cadmus` is currently limited to PubMed-based search queries. Similar queries can be generated through other publication databases and could be incorporated over time. Each database has its own syntax, which would require a conversion script and de-duplication steps to be added. `cadmus` has been designed for biomedical text retrieval, and performance has not been measured for searches outside of this domain. In other studies (20, 21), we re-processed XML and HTML files retrieved by `cadmus` using Auto-CORPus (22) and the Information Artifact Ontology (IAO) (23) to restore document hierarchy and focus the analysis on abstracts and result sections of articles. Future development will focus on expanding the codebase to support additional parsing strategies and output formats. This tool has been developed in Scotland, which is protected under UK law and backed by a commitment by Science, Technical and Medical Publishers to facilitate text and data mining studies (24) applicable to the UK and all areas of the European Union. Researchers in other legal jurisdictions may be vulnerable to copyright challenge and should adhere to any national guidance on text and data mining.

## Conclusion

`cadmus` provides a robust solution for large-scale biomedical full-text retrieval, addressing a challenge in biomedical text mining. By integrating multiple APIs, supporting diverse file formats, and offering efficient parsing and storage, it simplifies the creation of domain-specific corpora while maintaining high fidelity to the extracted scientific content. During our analysis, we demonstrate its ability to substantially increase biomedical concept coverage compared to abstracts alone. `cadmus` is freely available under an open-source licence, allowing anyone to create their biomedical domain-specific corpora for their text mining needs.

## AUTHOR CONTRIBUTIONS

Conceptualisation: IS; Methodology: JC, AL, and IS; Software: JC, and AL; Validation: JC, AL, and IS; Formal analysis: JC, AL, and IS; Investigation: JC, AL, and IS Resources: IS; Data Curation: JC, AL, and IS; Writing - Original Draft: JC, AL, and IS; Writing - Review & Editing: JC, AL, and IS; Visualisation: JC, AL, and IS; Supervision: IS. All authors have read and agreed to the published version of the manuscript.

